# CD163 protects against pulmonary injury and inflammation induced by acute O_3_ exposure

**DOI:** 10.64898/2026.06.01.726922

**Authors:** Samuel J. Cochran, Brett Saunders, Evangeline Schott, Katelyn Dunigan-Russell, Grace M. Hutton, Aaron Vose, Anastasiya Birukova, Cameron Rankin, Timothy J. McMahon, Hongmei Zhu, Valery V. Khramtsov, Murugesan Velayutham, Salik Hussain, Robert M. Tighe, Kymberly M. Gowdy

## Abstract

Ozone (O_3_)-driven pulmonary inflammation is partly regulated by damage associated molecular patterns (DAMPs) binding to scavenging receptors (SRs). However, how SRs and DAMPs regulate O_3_-induced pulmonary inflammation remains incompletely understood. CD163 is a SR responsible for clearing cell free hemoglobin (CFH), a DAMP which accumulates during acute pulmonary injury and is associated with worsening respiratory outcomes. We hypothesized that increased CD163 is necessary for reducing CFH levels and resolving O_3_-induced pulmonary injury. To test this hypothesis, we defined CD163 and CFH responses to O_3_ exposure in C57BL/6N (WT) and CD163 deficient (*Cd163*^*-/-*^) mice, as well as in human bronchoalveolar lavage fluid (BALF). In WT mice, lung *Cd163* expression was significantly increased by O_3_ during peak inflammation and declined 24 hours post exposure. Human exposure studies revealed a diversity of *Cd163* expression and a reduction of CFH following O_3_ exposure, suggesting regulation of this pathway in humans. When compared to WT mice, *Cd163*^*-/-*^ mice had augmented O_3_-induced pulmonary injury, inflammation, and oxidative stress. Further, the antioxidant EUK-134 did not reduce O_3_-induced pulmonary oxidative stress in *Cd163*^*-/-*^ mice, suggesting a role for CD163 in the pulmonary response to oxidative insults. Furthermore, compared to WT controls, *Cd163*^*-/-*^ mice receiving an oropharyngeal aspiration of CFH had a significant increase in airspace inflammation. Combined, these findings suggest that CD163 mediated clearance of CFH is involved in resolving O_3_-induced pulmonary injury, inflammation, and oxidative stress.

**New & Noteworthy:** Ozone (O_3_) is known to induce damage associated molecular patterns (DAMPs) which drive lung inflammation. The scavenging receptor, CD163, binds and clears the DAMP cell free hemoglobin (CFH), which accumulates during sterile lung injury. Our findings indicate that O_3_ exposure alters CD163 expression in the lung and that mice lacking *Cd163* expression have more lung inflammation. Our data indicate that CD163 serves a protective role in response to acute O_3_ exposure perhaps through CFH clearance.

## Introduction

Exposure to air pollution is a significant health concern, contributing to over 6.5 million premature deaths globally in 2015 (1, 2). In the United States, combustion emissions, including ultrafine particulate matter (PM_2.5_) and ozone (O_3_), contribute to an estimated 90,000 – 360,000 premature deaths annually (3-5). Ground level O_3_ forms in the lower atmosphere through a series of photochemical reactions involving heat, ultraviolet radiation, and primary air pollutants such as volatile organic compounds (VOCs) and nitrogen dioxide (NO_2_). Despite regulations set by the National Ambient Air Quality Standards (NAAQS), exposure to excess O_3_ concentrations between 2000 and 2002 resulted in an estimated average of 800 premature deaths and $5.7 billion in direct and indirect costs associated with healthcare access or lost productivity (6). Moreover, ground-level concentrations of O_3_ are projected to increase nationwide in the USA with warming ambient temperatures (7), making O_3_ exposure an increasingly salient concern for public health.

O_3_ exposure increases mortality and morbidity of cardiopulmonary diseases (8-11) in part through induction of lung inflammation. Upon inhalation, O_3_ reacts with molecules in the airway surface lining (ASL), forming lipid oxidation/ozonation products (LOP), protein carbonyls, and damage-associated molecular patterns (DAMPs) (12, 13). These DAMPs are recognized by pattern recognition receptors (PRRs), which initiate pro-inflammatory signaling cascades (14-16) that if unchecked can drive persistent tissue injury and organ dysfunction. One class of PRR, scavenger receptors (SRs), has been shown to regulate O_3_-induced pulmonary injury. Some SRs augment pulmonary inflammation (CD36), while others are protective (SR-A and MARCO) (13, 17). This suggests non-redundancy for SRs in the pulmonary immune response; however, their complex role is still not fully understood.

CD163 is one of the SRs that has been understudied in the context of air pollution-mediated lung injury. CD163 is a class B SR and is expressed on the surface of monocytes and macrophages (18). It is essential in pathogenic conditions where red blood cell (RBC) lysis generates cell free hemoglobin (Hb) (CFH) and heme (19). CFH can be cytotoxic when it accumulates, necessitating its clearance to limit organ injury and damage. Once CFH binds to the scavenger protein, haptoglobin (Hp) (18), the Hb-Hp complex is recognized and internalized by membrane-bound CD163 for degradation by heme oxygenase 1 (HO-1). This degradation generates anti-inflammatory degradation products: bilirubin and carbon monoxide (CO). The iron liberated through this process can then be exported from the cell or safely sequestered in intracellular ferritin complexes to prevent cytotoxic Fenton reactions from occurring (20, 21). Internalization of CD163 and metabolism of Hb-Hp complexes initiates a macrophage signaling cascade resulting in the production of the anti-inflammatory cytokine, interleukin-10 (IL-10) (22, 23). In addition to its membrane-bound activities, CD163 is also secreted in a soluble form (sCD163) during inflammation (24). Because of this, plasma sCD163 levels have been used as a disease activity biomarker in diabetes, obesity, and cardiovascular diseases (25-28). Despite the understanding of CFH and CD163 in immune responses, the functional relevance of CD163 and sCD163 in scavenging CFH and promoting anti-inflammatory responses in air pollution-induced lung injury is not currently understood. Therefore, we hypothesized that CD163 is necessary for the upregulation of anti-inflammatory pathways involved in resolution of pulmonary inflammation caused by acute O_3_ inhalation. In this study, the loss of CD163 resulted in augmented O_3_-induced pulmonary injury, inflammation, and oxidative stress. Whole-body *Cd163* knockout mice had augmented pulmonary inflammation in response to pulmonary CFH, suggesting a mediating role of CFH in O_3_-induced inflammation and injury.

## Methods

### Experimental animals

Female, wild type (WT), C57BL/6N mice were purchased from Taconic (Taconic Biosciences, NY, USA). *Cd163*^tm1(KOMP)Vlcg^ (*Cd163*^*-/-*^) whole body knockout (KO) mice were a generous gift from Stewart Levine, MD (National Heart, Lung, and Blood Institute, Bethesda, MD), originally procured from the Knockout Mouse Project (KOMP) (29). All mice were housed in the Ohio State University animal facilities with support from University Laboratory Animal Resources. Mice were maintained at 22.2 °C ± 2.2 °C, on a 12 hour light and dark cycle, with ad libitum access to food (Teklad 7912, Envigo, IN, USA) and water (Lixit, CA, USA). *Cd163*^*-/-*^ mice genotypes were confirmed as previously described (30). All experiments were performed in accordance with the Animal Welfare Act and the United States Public Health Service Policy on Humane Care and Use of Laboratory Animals after review by the Animal Care and Use Committee of Ohio State University.

### Murine in vivo exposure studies

Mice were exposed to either filtered air (FA) or 1 ppm O_3_ for 3 hours, as previously described (31, 32). This dose is directly proportional to human O_3_ exposures on an ‘O_3_ action day’ (33, 34). Temperature and humidity were monitored continuously. In a separate set of experiments, a subset of mice received a single intraperitoneal injection (200 μL) of the antioxidant, EUK-134 (Millipore Sigma, MA, USA), 1 hour before exposure as previously described (35). EUK-134 is a synthetic superoxide dismutase and catalase mimetic, thereby effectively reducing reactive oxygen species (ROS) commonly generated following O_3_ inhalation (35). All experiments involving cell free hemoglobin (CFH) were performed as previously described (36). Briefly, mice were anaesthetized with isoflurane and then received an oropharyngeal (o.p.) aspiration of 1 mg/mL native human CFH (100 μL; Cat. No. CSI9668B Lot No. 4114204, Cell Sciences, MA, USA) suspended in sterile PBS. This biologically relevant dose is based on an average CFH concentration measurable in undiluted pulmonary edema fluid from patients with acute respiratory distress syndrome (ARDS) (37). For all experiments, mice were euthanized via a single intraperitoneal injection (200 μL) of a ketamine (100 mg/kg) and xylazine (10 mg/kg) mixture in sterile saline (38).

### Bronchoalveolar lavage fluid (BALF) collection and analysis

The right lung was lavaged three times with 26.25 mL/kg of 1X Dulbecco’s phosphate buffered saline (PBS; Life Technologies, NY, USA). The BALF was centrifuged at 460 x g for 6 minutes at 4 °C. The BALF supernatant was used to measure albumin (Immunology Consultants Laboratory, OR, USA) and total protein via bicinchoninic acid (BCA) Protein Assay Kit (Thermo Fisher Scientific, MA, USA). To the cell pellet, 1 mL of red blood cell lysis buffer (in distilled H_2_O: 168.3 μM NH_4_CL, 9.99 μM KHCO_3_, 0.125 μM EDTA) was added, vortexed, and incubated for 1 min at room temperature. To stop the reaction, 1X Dulbecco’s phosphate buffered saline (PBS; Life Technologies, NY, USA) was added. The airspace cells were again centrifuged 460 x g for 6 minutes at 4 °C, then resuspended in 1 mL of 10% heat inactivated fetal bovine serum (Life Technologies, NY, USA) in PBS. The immune cells from the BALF cell pellet were quantified via Bright-line hemocytometer (Hausser Scientific, Horsham, PA) and identified using Cytospin 4 (Thermo Fisher Scientific, MA, USA) and Kwik-Diff™ Kit solution (Epredia, MI, USA). Concentrations of select cytokines and chemokines were measured via multiplex ELISA (Meso Scale Diagnostics, MD, USA).

### Human exposure studies and BALF collection

Cell pellets and cell-free supernatants were obtained from BAL performed on healthy human participants exposed to FA and O_3_, as previously described (39). Participants were recruited at Duke University in two Duke Institutional Review Board approved clinical studies (Pro00088966 and Pro00100375). Participants were enrolled after completion of informed consent. Demographic information is listed in Table 1. Participants were exposed to FA or 0.200 ppm +/- 0.015 ppm O_3_ for 2.25 hours during which time participants alternated rest and walking on a treadmill at ∼2-3 mph to simulate mild exertion. The O_3_ dose exceeds NAAQS (80 ppb for 8-hour average) but is comparable to ground level concentrations during ‘ozone action days’ in the Research Triangle area. Temperature and relative humidity during exposures ranged from 20-23 °C and 45-55%, respectively. The exposures were performed in a blinded, case-crossover study design where participants were randomly assigned to FA or O_3_ exposure and then exposed to the alternative condition after a minimum 21-day washout period. Thus, each participant served as their own control. Approximately 21 hours after each exposure, participants underwent a flexible bronchoscopy with BAL. A flexible bronchoscope was wedged into the right middle lobe and aliquots of 30 mL to a total of 150 mL were discharged into the lungs and removed with gentle syringe suction to obtain BALF.

**Table 1.**
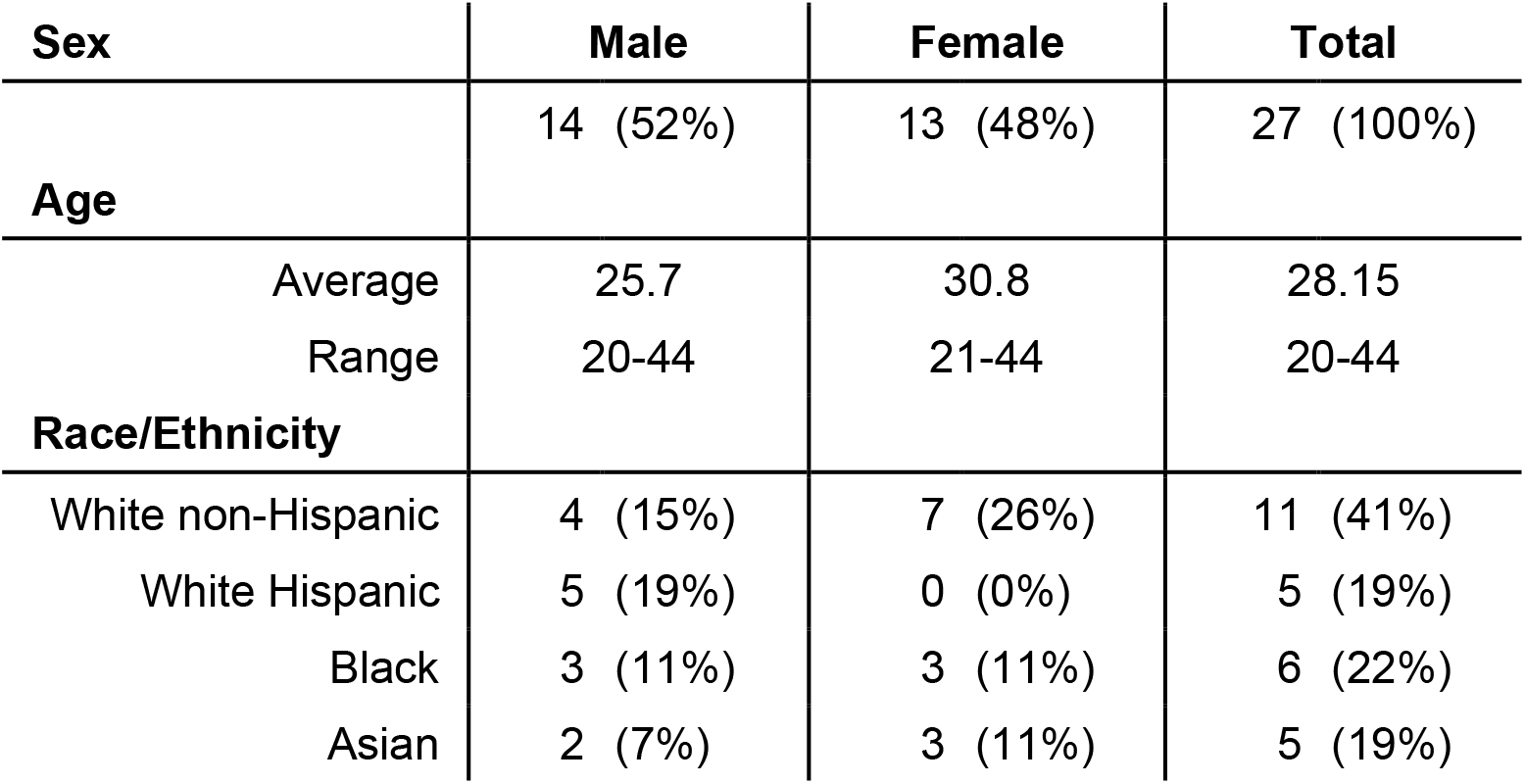
Demographic information of recruited human participants.

| **Sex** | **Male** | **Female** | **Total** |
| --- | --- | --- | --- |
|  | 14 (52%) | 13 (48%) | 27 (100%) |
| **Age** |  |  |  |
| Average | 25.7 | 30.8 | 28.15 |
| Range | 20-44 | 21-44 | 20-44 |
| **Race/Ethnicity** |  |  |  |
| White non-Hispanic | 4 (15%) | 7 (26%) | 11 (41%) |
| White Hispanic | 5 (19%) | 0 (0%) | 5 (19%) |
| Black | 3 (11%) | 3 (11%) | 6 (22%) |
| Asian | 2 (7%) | 3 (11%) | 5 (19%) |

### Analysis of airspace hemoglobin and hemoglobin-associated proteins

Human and murine BALF hemoglobin (Hb) was measured using spectroscopy, as previously described (40-42). Murine BALF haptoglobin (Hp) was measured by ELISA (ab15714, Abcam, Cambridge, UK) and murine BALF hemopexin (Hx) was measured by ELISA (ab157716, Abcam, Cambridge, UK). Soluble CD163 (sCD163) in human BALF was measured via ELISA (DC1630 Bio-Techne, MN, USA).

### RNA isolation and quantitative polymerase chain reaction (qPCR)

Murine left lungs were collected, snap-frozen in liquid nitrogen, and stored at −80 degrees Celsius. The frozen tissue was placed into 2 mL tubes containing 1 mL of tissue lysis buffer (1% β-mercaptoethanol in RLT Buffer, RNeasy MiniKit, Qiagen, Hilden, Germany) and a 5 mm steel bead. The tissue was homogenized at 50 oscillations per second for 10 minutes (TissueLyser LT, Qiagen, Hilden, Germany) while at 4 °C, then centrifuged at 10,000 x g for 5 minutes while at room temperature. The supernatants were collected and processed according to instructions in the Qiagen RNeasy MiniKit. Isolated RNA was quantified using a SpectraDrop Micro-Volume Microplate and a SpectraMax iD3 (Molecular Devices, CA, USA). cDNA was synthesized from RNA using the High-Capacity cDNA Reverse Transcription Kit (Thermo Fisher Scientific, MA, USA) and a ProFlex PCR System (Thermo Fisher Scientific, MA, USA). The resulting cDNA was diluted to a 5X volume using nuclease free water (Ambion, TX, USA) and store at −20 degrees Celsius. Relative expression of select genes was measured using Taqman assays: heme oxygenase 1, *Hmox1* (Mm00516005_m1); interleukin-10, *Il10* (Mm01288386_m1); and toll-like receptor 4, *Tlr4* (Mm00443258_m1, Thermo Fisher Scientific, MA, USA). All assays were performed using Taqman PCR Mix (Applied Biosystems, MA, USA) on the StepOnePlus Real-Time PCR System (Applied Biosystems, MA, USA). Fold change expression of mRNA was calculated using the 2^-ΔΔCT^ method and normalized relative to *18S* gene expression (Mm03928990_g1) as previously described (43). Human *Cd163* expression was measured from BALF cells using SYBR Assay (Thermo Fisher Scientific, MA, USA) and was normalized to human 18S and then normalized to the filtered air control to the individual subject.

### Measurement of airspace oxidative stress

Oxidizing potential of bronchoalveolar lavage and serum samples were measured by electron paramagnetic resonance (EPR) spectroscopic technique using 1-hydroxy-3-carboxymethyl-2,2,5,5-tetramethyl-pyrrolidine (CMH) spin probe (Enzo Life Science, Long Island, NY). Sample collection and preparation was performed following our previously described methodology (44). EPR spectra were recorded using a Bruker ELEXSYS E580 spectrometer (Bruker BioSciences, Billerica, MA, USA) operating at X-band with 100 kHz modulation frequency. The following EPR instrument settings were used, microwave frequency, 9.854 GHz; center field, 3495 G; sweep width, 100 G; microwave power, 23.77 mW; modulation amplitude, 1 G; modulation frequency, 100 kHz; receiver gain, 60 dB; conversion time, 29.3 ms, sweep time, 30 s; number of scans, 1. EPR data acquisition was performed using Bruker Xepr software. Airspace oxidative stress was also measured by OxiSelect™ ELISA (Fisher Scientific, Waltham, MA) detection of protein carbonyl levels in BALF.

### Statistical analyses

All statistical analyses were performed in GraphPad Prism version 10.0.0 for Windows (GraphPad Software, MA, USA). Results are reported as averages with standard error of the mean (SEM) represented in the error bars. Kruskal-Wallis with Dunn’s test adjustment for multiple comparisons was performed when comparing conditions (e.g. timepoints) to baseline (Figure 1). When assumptions of normality were not met for Two-way ANOVA, Multiple Mann-Whitney U-tests were instead performed to compare significant differences between groups across two separate independent variables. Holm’s post hoc adjustment for multiple comparisons was used to control the type one error rate in these three kinds of tests. Paired Wilcoxon Ranked Sign tests were used to analyze changes in human participants between the two exposure conditions. A p-value < 0.05 was considered significant for all tests. Adjusted p-values are reported when relevant.

**Figure 1.**
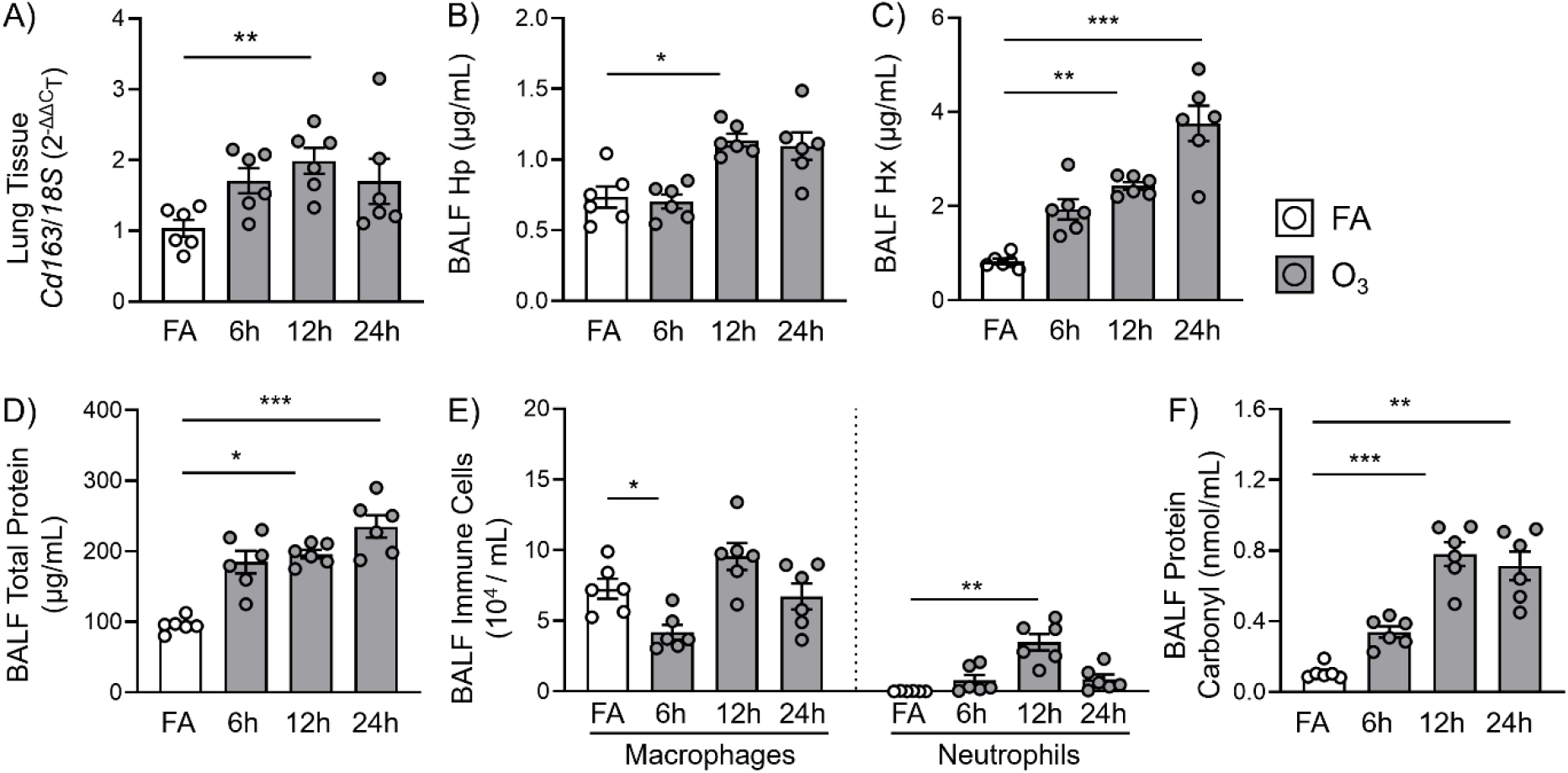
O_3_ increases pulmonary Cd163 expression and heme/hemoglobin scavenging protein concentrations. Female, C57BL/6N wildtype (WT) mice, aged 11 weeks old were exposed to either filtered air (FA) or 1 ppm ozone (O_3_) for 3 hours. Mice were euthanized 6, 12, and 24 hours after exposure to collect lung tissue and lung bronchoalveolar lavage fluid (BALF). A) Expression of Cd163 was measured via rtPCR in homogenized lung tissue. B-C) BALF concentrations of haptoglobin (Hp) and hemopexin (Hx) were measured by ELISA. D) Airspace lung injury was measured by concentration of total protein in the BALF. E) Lung inflammation was measured by macrophage numbers and neutrophil infiltration in the BALF. F) Oxidative stress was measured by concentration of BALF protein carbonyl concentration. Kruskal-Wallis with Dunn’s test for multiple comparisons post-hoc; n = 6 /group; *p<0.05; **p<0.01; ***p<0.001

## Results

### Acute O_3_ exposure increases pulmonary expression of Cd163 which is associated with increased airspace levels of heme/hemoglobin scavenging proteins

To determine whether CD163 mediates the pulmonary response to O_3_, *Cd163* expression was measured in homogenized lung tissue and macrophages collected from BALF of mice exposed to FA and O_3_. *Cd163* expression was significantly increased 12 hours after exposure in lung tissue (Figure 1A). Airspace concentration of the Hb-scavenging protein, haptoglobin (Hp), also peaks at 12 hours (Figure 1B), mirroring significant increases in airspace concentrations of the heme-scavenging protein, hemopexin (Hx) (Figure 1C). These measures coincided with O_3_ driven increases in BALF total protein, at 12 and 24 hours after exposure (Figure 1D). Additionally, O_3_ driven lung inflammation occurred 12 hours after exposure, as evidenced by significant increases in numbers of BALF neutrophils as well as a non-significant increase in BALF macrophages (Figure 1E). This coincided with a 12-hour peak in airspace oxidative stress, as measured by BALF protein carbonyl concentration (Figure 1F). Combined, these data indicate that O_3_ induces lung injury, inflammation, and oxidative stress concomitant with upregulation of *Cd163* expression and production of heme and hemoglobin scavenging proteins.

### Acute O_3_ exposure alters airspace CD163 and hemoglobin levels in humans

To determine if our findings in mice translate to human O_3_ exposure, we assessed *Cd163* expression, sCD163 concentrations, and CFH in human BALF from young, healthy humans. Participants without evidence of respiratory disease who were exposed to acute FA and O_3_ in a case-crossover study design. O_3_ exposure, when normalized to the FA exposure in the individual participant, led to an average, though non-statistical, increase in *Cd163* mRNA expression in BALF cells (Figure 2A). Interestingly, O_3_ exposure resulted in a significant decrease in the airspace concentration of CFH (Figure 2B) and no change in airspace sCD163 (Figure 2C).

**Figure 2.**
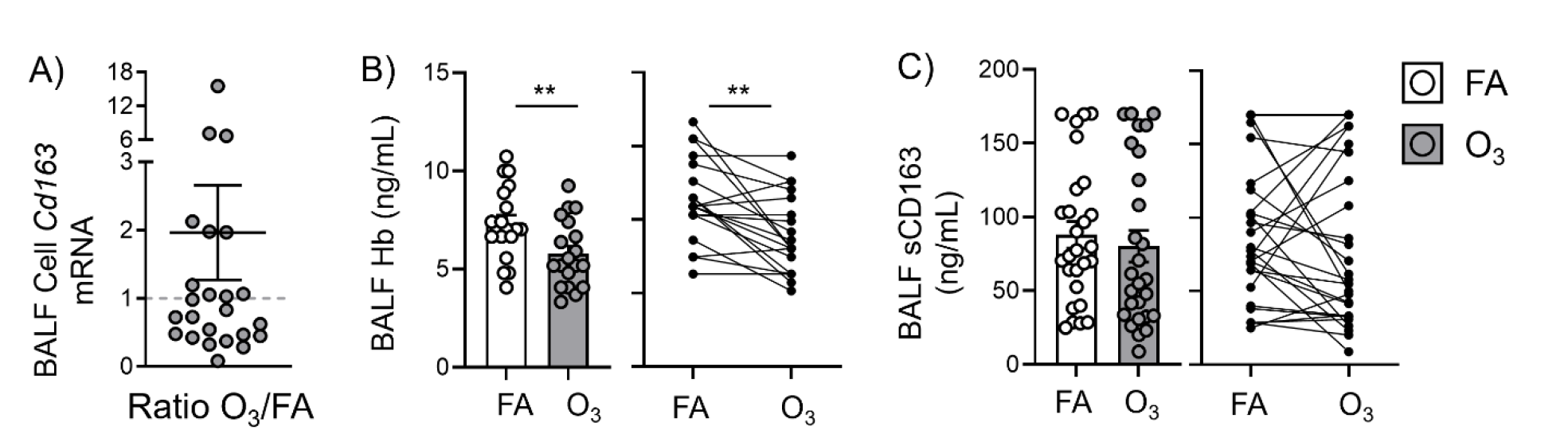
O_3_ increases human airspace cell Cd163 expression and decreases airspace Hb and sCD163 concentrations. In a random, blinded case-crossover study design, human participants (male and female, ages 18-35) were exposed to filtered air (FA) and ozone (O_3_) (0.200 ppm) for 2.25 hours, then exposed to alternative condition after a 21-day washout period. Bronchoalveolar lavage fluid (BALF) was collected 18 – 24 hours following exposures. A) Cd163 expression was measured via rtPCR in the BALF cell pellet. B) BALF hemoglobin (Hb) was measured by spectrometry. C) BALF soluble CD163 (sCD163) was measured by ELISA. Paired, Wilcoxon Ranked-Sign test; n = 17 – 27 /group; **p<0.01

### CD163 protects against O_3_-induced pulmonary injury and inflammation

To better understand whether O_3_-induced lung injury and inflammation is mediated by CD163, lung tissue and BALF were collected from whole body *Cd163*^*-/-*^ and WT mice 12 hours and 24 hours following acute O_3_ exposure. Compared to FA controls, WT and *Cd163*^*-/-*^ mice both exhibited significantly increased BALF total protein 12 hours after O_3_ exposure (Figure 3A). This increase was augmented in *Cd163*^*-/-*^ compared to WT animals, indicating a protective role of CD163 against O_3_-induced lung injury. Following this trend, lung inflammation was also augmented in *Cd163*^*-/-*^ compared to WT mice 12 hours after O_3_ exposure, as indicated by significantly increased BALF total immune cells; significantly increased BALF macrophages; and a trending increase in BALF neutrophils (Figure 3B). Concentrations of BALF cytokines/chemokines 12 hours following O_3_ were not significantly different between WT and *Cd163*^*-/-*^ mice (Supplemental Figure 1). In contrast to the 12-hour timepoint, BALF total protein was not augmented in *Cd163*^*-/-*^ mice 24 hours after O_3_ exposure (Figure 3C), suggesting that 12 hours is a critical time point in CD163 response to lung injury. Still, lung inflammation was augmented in *Cd163*^*-/-*^ mice 24 hours after O_3_ exposure, indicated by significantly increased neutrophilia in the BALF (Figure 3D). This coincided with augmented oxidative stress in *Cd163*^*-/-*^ compared to WT mice, as measured by increased BALF oxidants (Figure 3E).

**Figure 3.**
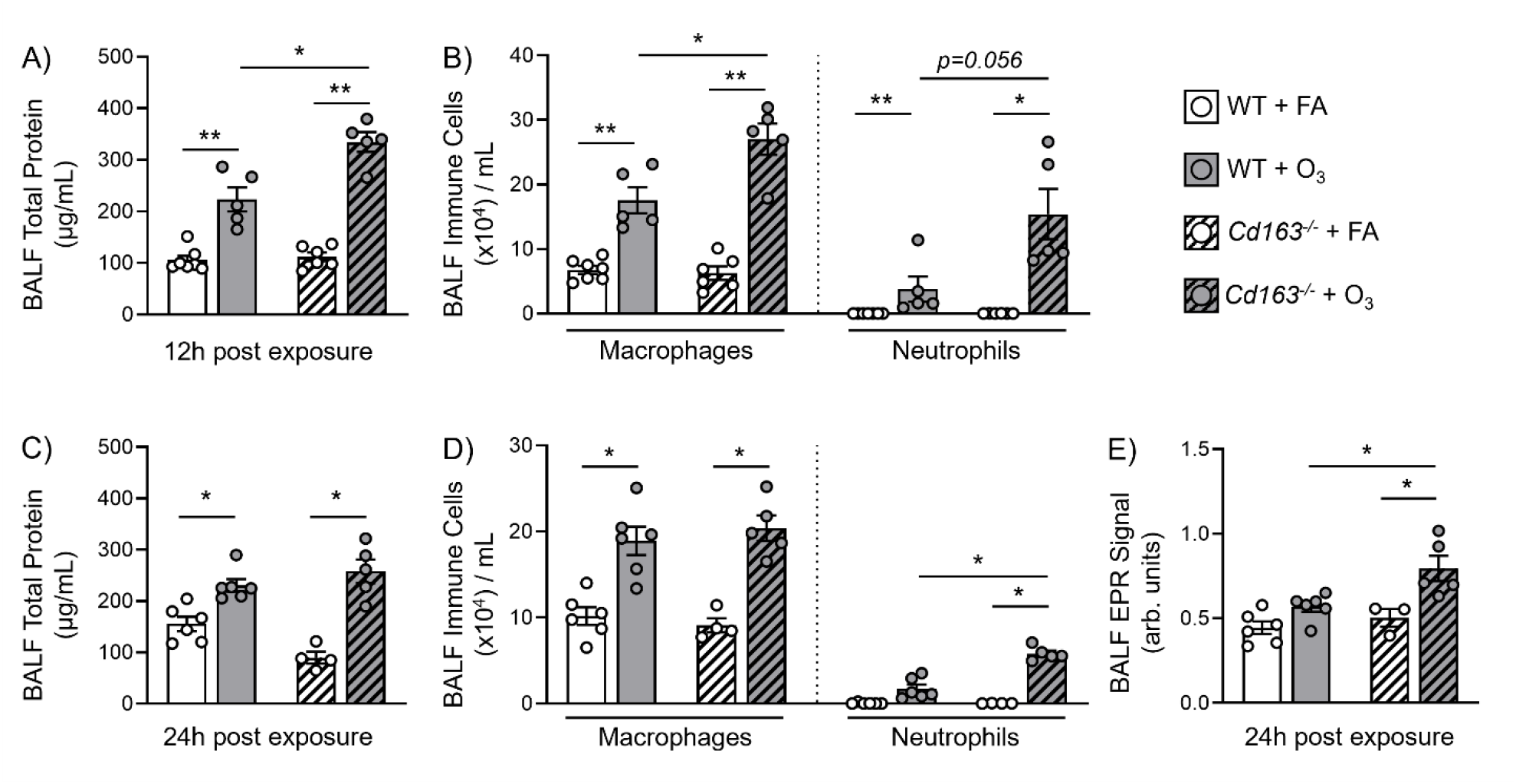
O_3_-induced lung injury, inflammation, and oxidative stress is augmented in Cd163^-/-^ mice. Female, C57BL/6N wildtype (WT) and Cd163^-/-^ mice, aged 8 – 12 weeks old were exposed to either filtered air (FA) or 1 ppm ozone (O_3_) for 3 hours. Mice were euthanized 12 and 24 hours after exposure to collect lung bronchoalveolar lavage fluid (BALF). Airspace lung injury was measured by concentration of airspace total protein in the BALF, 12 hours (A) and 24 hours (C) after exposure. Lung inflammation was measured by macrophage numbers and neutrophil infiltration in the BALF, 12 hours (B) and 24 hours (D) after exposure. E) Airspace reactive oxygen species were measured by electron paramagnetic resonance spectroscopy (EPR, arbitrary units), 24 hours after exposure. Mann-Whitney U Test with Holm’s adjustment for multiple comparison; n = 4 – 6 /group; *p<0.05; **p<0.01

### CD163 mediates O_3_-induced pulmonary oxidative stress

To better understand the role of CD163 in mediating oxidative stress in the lung and to identify potential therapies, we attempted to reduce the augmented oxidative stress in *Cd163*^*-/-*^ mice by administering a prophylactic dose of EUK-134. EUK-134 is a synthetic superoxide dismutase and catalase mimetic, which preferentially targets two types of ROS (O_2_^-^ and H_2_O_2_) commonly produced in the lung following O_3_ inhalation (35). Although EUK-134 was protective against O_3_-induced oxidative stress in WT mice, by comparison, oxidative stress was significantly increased in *CD163*^*-/-*^ mice receiving EUK-134 (Figure 4A). Additionally, EUK-134 treatment had no significant effect on lung BALF total protein or neutrophilia in either *Cd163*^*-/-*^ or WT mice (Figure 4B and 4C). These findings indicate reduced efficacy for EUK-134 in mediating O_3_-induced oxidative stress in mice lacking *Cd163*.

**Figure 4.**
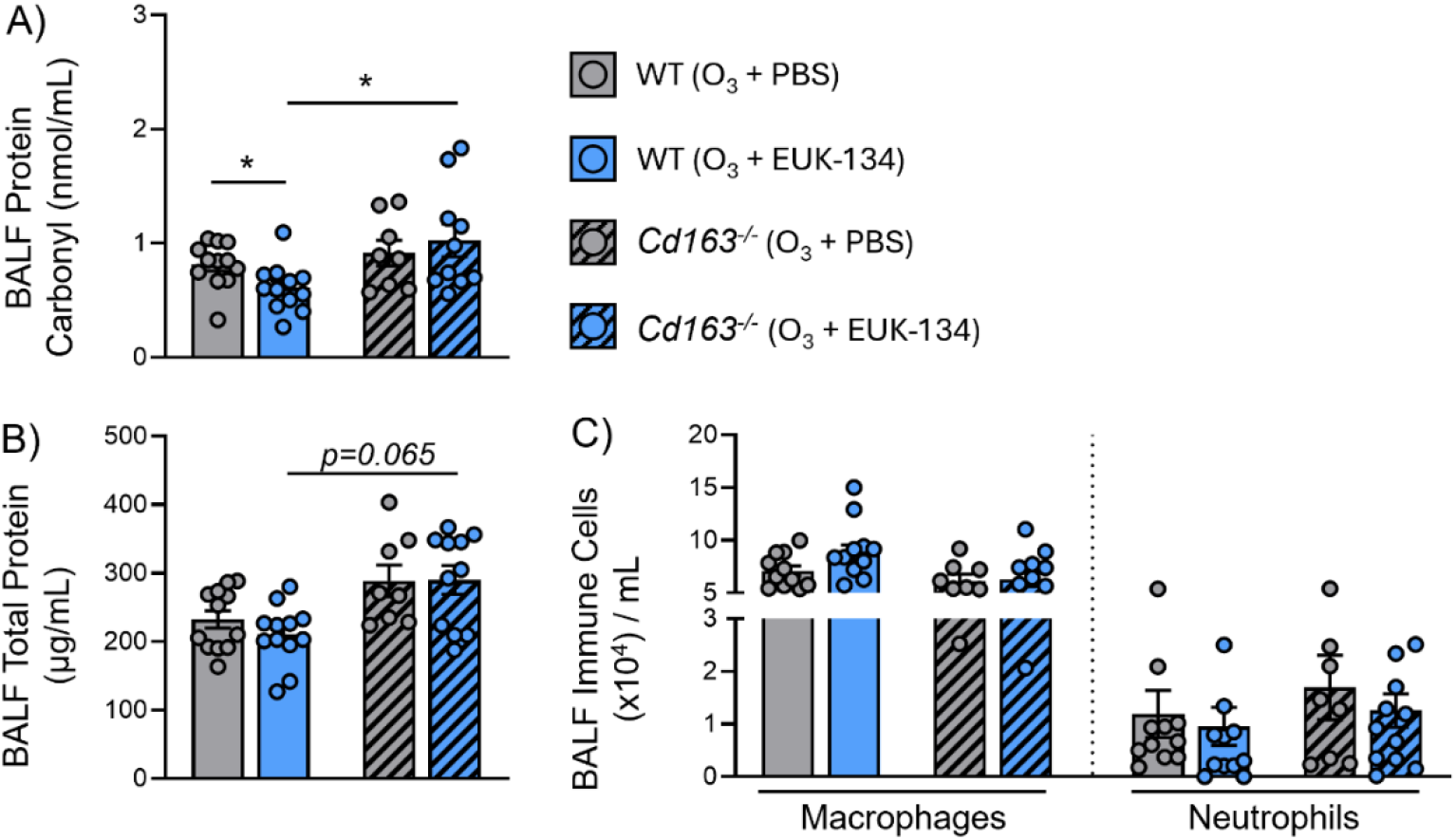
Antioxidant (EUK-134) administration does not rescue augmented oxidative stress in Cd163^-/-^ mice. Female, C57BL/6N wildtype (WT) and Cd163^-/-^ mice, aged 8 – 12 weeks old were exposed to 1 ppm ozone (O_3_) for 3 hours. Mice received an intraperitoneal (i.p.) injection of the superoxide dismutase (SOD) and catalase mimetic, EUK-134 (10 mg/kg), or vehicle control (0.9% sterile saline) approximately 1 hour prior to exposure. Mice were euthanized 24 hours after exposure to collect lung bronchoalveolar lavage fluid (BALF). A) Airspace oxidative stress was concentration of BALF protein carbonyl. B) Airspace lung injury was measured by concentration of airspace total protein in the BALF. C) Lung inflammation was measured by macrophage numbers and neutrophil infiltration in the BALF. Mann-Whitney U Test with Holm’s adjustment for multiple comparison; n = 11 – 12 /group; *p<0.05

### CD163 expression protects against CFH-induced lung inflammation

We predicted that impaired CFH clearance drives the increased oxidative stress, lung injury, and inflammation in mice lacking *Cd163*. To determine whether CFH clearance is inhibited in *Cd163*^*-/-*^ mice, lung tissue and BALF were collected from *Cd163*^*-/-*^ and WT mice 24 hours following an oropharyngeal instillation of CFH (1 mg/mL). This biologically relevant dose is based on an average CFH concentration measurable in undiluted pulmonary edema fluid from patients with ARDS (37) and this timepoint corresponds to previously published peaks in CFH-induced lung inflammation (36). Administered CFH did not increase BALF total protein in either WT or *Cd163*^*-/-*^ mice (Figure 5A). Though BALF macrophage numbers were mostly unchanged, CFH administration led to an increase in neutrophil infiltration in both groups (Figure 5B). Airspace neutrophilia was further augmented in *Cd163*^*-/-*^ mice compared to WT, indicating increased airspace inflammation perhaps due to impaired CFH clearance (Figure 5B). Finally, we measured airspace oxidative stress by measuring BALF protein carbonyl levels. Unlike findings following O_3_ exposure (Figure 2E), we did not detect augmented oxidative stress in *Cd163*^*-/-*^ compared to WT mice (Figure 5C). We confirmed that BALF concentrations of CFH were not augmented in *Cd163*^*-/-*^ mice. This could indicate compensatory mechanisms of CFH or heme uptake. However, the increase in detectable CFH after CFH administration was only significant in the *Cd163*^*-/-*^ group (Figure 5D) and BALF Hp concentrations were augmented in the *Cd163*^*-/-*^ mice compared to the CFH-treated WT (Figure 5E). Given these trends, we assessed changes in transcription of genes linked to CD163 internalization. *Hmox1* (antioxidant) and *Il10* (anti-inflammatory) expression were both elevated among WT and *Cd163*^*-/-*^ mice dosed with CFH (Supplementary Figure 2A and B). There were no trends with expression of *Tlr4* (Supplementary Figure 2C), which is known to upregulate pro-inflammatory pathways when binding CFH (45). Taken together, our data indicate that *Cd163*^*-/-*^ mice dosed with CFH have an increase in airspace neutrophils, hemoglobin and haptoglobin, suggesting potential defects in CFH clearance.

**Figure 5.**
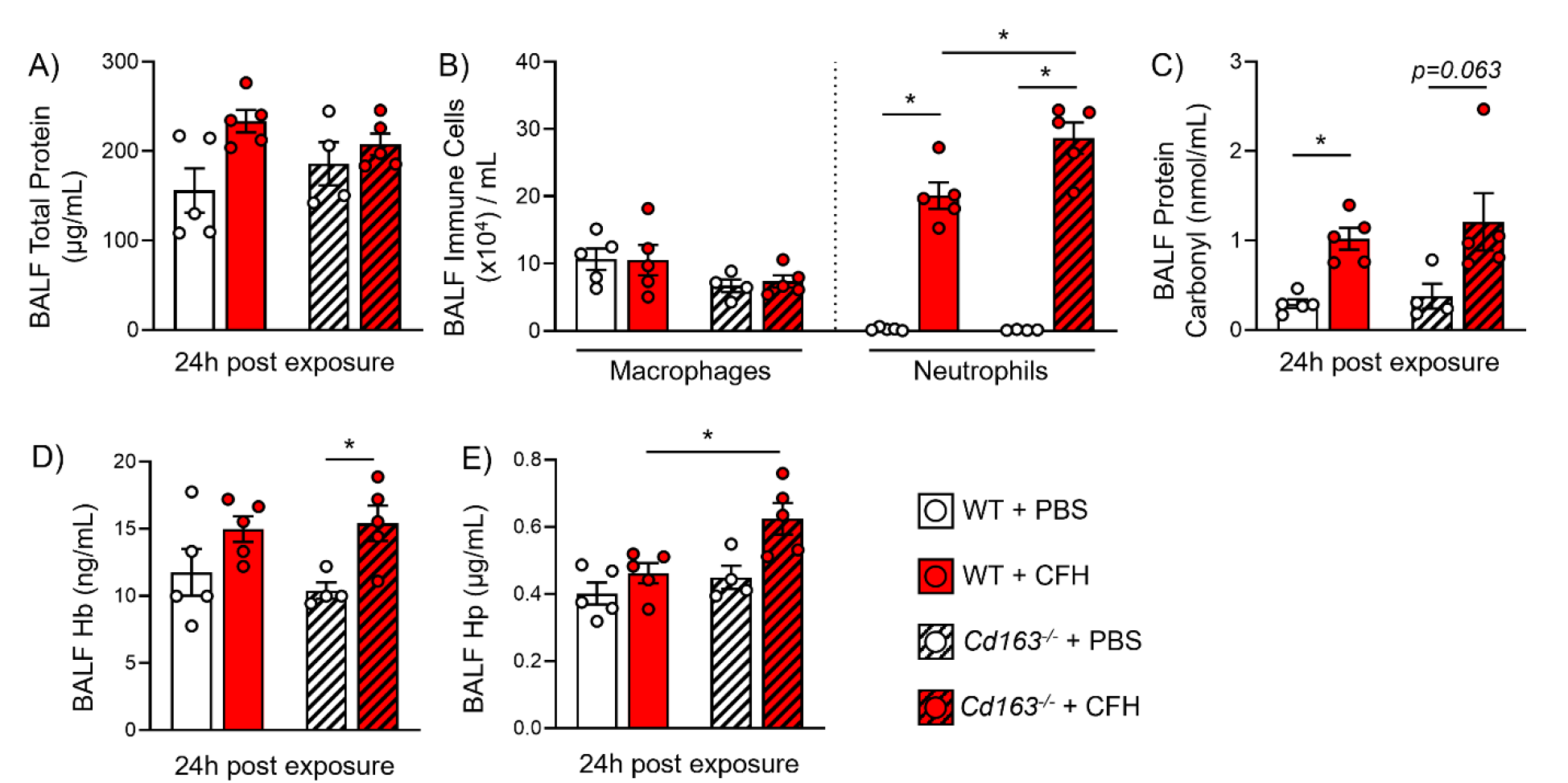
CFH oropharyngeal aspiration increases neutrophilia and augments oxidative stress in Cd163^-/-^ mice. Female, C57BL/6N wildtype (WT) and Cd163^-/-^ mice, aged 8 – 12 weeks old were dosed with 100 μL of either sterile PBS or 1 mg/mL CFH oropharyngeally (o.p.). Mice were euthanized 24 hours after exposure to collect bronchoalveolar lavage fluid (BALF). A) Airspace lung injury was measured by concentration of airspace total protein in the BALF. B) Lung inflammation was measured by macrophage numbers and neutrophil infiltration. C) Oxidative stress was measured by concentration of BALF protein carbonyl. D) hemoglobin (Hb) was measured by spectrometry. E) Haptoglobin (Hp) production was measured by ELISA. Mann-Whitney U Test with Holm’s adjustment for multiple comparison; n = 4 – 6 /group; *p<0.05

## Discussion

Different SRs can augment or attenuate inflammatory responses during sterile lung injury (13, 17, 46). However, the function of many SRs remains poorly understood, especially in the context of O_3_-induced lung injury. Here, we investigated whether CD163 facilitates resolution of O_3_-driven lung injury and inflammation through its primary role in clearing CFH. Our data indicates that acute O_3_ inhalation increases *Cd163* expression in both humans and mice. Moreover, CD163 is partially protective against O_3_-induced markers of lung injury, inflammation, and oxidative stress. Interestingly, antioxidant administration was ineffective in reducing augmented oxidative stress in *Cd163*^*-/-*^ mice. Finally, we demonstrate that CD163 is protective in reducing inflammation induced by airspace CFH. Our study is the first to investigate CD163 in this novel context and the first to implicate CFH as a DAMP in O_3_-driven lung pathology. Cumulatively, our findings 1) indicate that CD163 reduces lung injury, inflammation, and oxidative stress caused by acute O_3_ inhalation and, 2) suggest that this mechanism occurs through clearance of O_3_-induced CFH accumulation in the lung.

The data presented in this study align with previous findings reported by our lab on other SR-facilitated DAMP clearance mechanisms. Our group has shown that another SR, SR-BI, is upregulated in the lungs following acute O_3_ exposure and that SR-BI functions to reduce O_3_-induced pulmonary inflammation by facilitating clearance of another class of DAMPs, oxidized phospholipids (46). Here, we present data demonstrating that acute O_3_ exposure upregulated expression of *Cd163* in murine whole lung tissues. This aligns with our previous findings that SRs canonically functions to clear DAMPs, which can induce oxidative stress and lead to augmented pulmonary pathologies (36, 47). In support of the concept of CFH clearance, we observed O_3_-induced *Cd163* upregulation concomitant with increases in BALF Hp and Hx concentrations. This suggests that along with increased *Cd163* there is an increase in the associated machinery required to sequester CFH, supporting its relevance in O_3_-induced pulmonary inflammation. Future studies could consider how increasing this clearance could be used to limit adverse health effects of ozone.

Although our initial studies in mice largely support our hypothesis, some incongruencies with our predictions required investigation of O_3_-induced *Cd163* expression in the human lung. Our findings in humans reproduced key findings in mice, with a few differences that could be explained by time point, gene expression between mice and humans, and inter-individual variability in human responses. As in mice, acute O_3_ inhalation increased average *Cd163* expression in the BALF cell pellet. However, this approximate doubling in *Cd163* expression was not statistically different, as there was significant inter-individual variation in response to O_3_. While some participants had reduced *Cd163* expression, several others had significantly upregulated *Cd163* expression, which may reflect the dichotomy between O_3_ responders and non-responders as has been well described (48, 49). Alternatively, this may reflect differences in the cell populations sampled between mice and humans. Prior data suggests that *Cd163* is highly expressed in murine interstitial macrophages (IMs) (30); however, IMs are not well sampled in human BALF. This could explain reduced *Cd163* expression in human BALF cells compared to murine lung tissue. Additionally, prior work has identified differences between rodent and human *Cd163* expression, namely human monocyte derived macrophages (MDM) produce CD163, unlike rodent MDM (18, 50). Future studies will need to consider cell type specific expression of CD163 in mice and humans to resolve the sources of increased CD163 following exposure. In addition to differences in CD163, we observed a significant decrease in human BALF Hb but no difference in BALF sCD163. This could be due to upregulation of other mechanisms involved in clearance of CFH and heme, such macropinocytosis (51), and uptake of heme-hemopexin complexes by CD91 (52). It is also possible that our findings result from BALF being collected approximately 24 hours after O_3_ exposure, at which point resolution of inflammation has been initiated (53). In fact, we observed similar trends in WT mice exposed to O_3_. Lung tissue and BALF macrophage expression of *Cd163* declines 24 hours post O_3_ exposure, concomitant with a plateauing of BALF Hp concentration. Thus, it possible that functional CD163 in human lungs leads to increased Hb clearance through sCD163 internalization. Further investigation at earlier timepoints could prove enlightening but may be prohibitive due to the difficulty associated with experiments using human participants.

To better understand the role of CD163 in mitigating O_3_-induced lung injury and inflammation, we exposed *Cd163*^*-/-*^ and WT mice to FA and O_3_ and compared responses at 12 and 24 hours post exposure. At 12 hours post O_3_ exposure, *Cd163*^*-/-*^ mice when compared to WT mice, had augmented BALF levels of total protein and neutrophilia, indicative of enhanced pulmonary injury and inflammation. At 24 hours post exposure, this pattern was sustained for neutrophilia but not BALF protein, suggesting that CD163 may be involved in early responses to injury or that compensatory responses begin to activate in *Cd163*^*-/-*^ mice at 24 hours (51, 54). There were augmented BALF protein carbonyl concentrations in *Cd163*^*-/-*^+O_3_ mice compared to WT+O_3_ mice, even at 24 hours, which suggests that CD163 may function through reduction of pulmonary oxidative stress. To investigate this line of inquiry, prior to O_3_ exposure, WT and *Cd163*^*-/-*^ mice received a prophylactic dose of EUK-134. EUK-134 is a mimetic for both SOD and catalase, thereby converting two important ROS (O_2_^-^ and H_2_O_2_) safely to H_2_O. Furthermore, it has been shown to be effective in reducing lung injury and inflammation caused by ultrafine carbon black (CB) and O_3_ inhalation co-exposure (35). In addition to different exposures (O_3_ vs CB+O_3_), other significant differences from the previous study include dose and sex of the animals. Potentially, greater benefit in CB+O_3_ exposure is due to a reduction in acellular oxidants generated on particle surfaces, which play a significant role in co-exposure toxicity as well as differential oxidant generation kinetics and nature of oxidants in O_3_ vs CB+O_3_ exposures. Although we did not see a reduction in BALF total protein or neutrophilia, EUK-134 did significantly reduce BALF protein carbonyls in WT mice exposed to O_3_. Interestingly, EUK-134 did not reduce oxidative stress in *Cd163*^*-/-*^ mice; in fact, BALF protein carbonyl levels were significantly increased in *Cd163*^*-/-*^+EUK-134 compared to WT+EUK-134 mice. One possible explanation for this response is that CD163 could be involved in antioxidant pathways targeting other ROS besides those targeted by EUK-134 (i.e. O_2_^-^ and H_2_O_2_). Further research into specific ROS types present in WT vs *Cd163*^*-/-*^ animals could elucidate this possibility and identify potential therapeutic targets for improving CD163-mediated antioxidant activity.

We next sought to determine whether the responses observed in *Cd163*^*-/-*^ mice are driven by the canonical role of CD163 in clearing CFH accumulation. CFH instillation resulted in non-significant increases in BALF total protein in both WT and *Cd163*^*-/-*^ groups, which could be confounded by the presence of CFH and associated proteins such as Hp and Hx. In fact, CFH induced augmented Hp concentrations in *Cd163*^*-/-*^ compared to WT BALF. This coincided with augmented neutrophilia in *Cd163*^*-/-*^+CFH mice compared to WT+CFH mice, providing further evidence that CD163 is important in mitigating the pro-inflammatory effects of this lung DAMP. It is also important to highlight that CFH drives increased airspace neutrophilia in both WT and *Cd163*^*-/-*^ mice. These data suggest that there are likely differences in the level of pulmonary CFH accumulation induced by O_3_ and direct exposure to a biologically relevant dose of CFH (36). Although inflammation was augmented in *Cd163*^*-/-*^+CFH mice, we did not observe the same trend for BALF protein carbonyl; in fact, there was only a trending increase. Thus, the canonical role of CD163 in mediating CFH clearance may not be sufficient to explain augmented oxidative stress in *Cd163*^*-/-*^ mice exposed to O_3_. There is evidence for other pathways involved in CFH uptake, which could be compensatory in *Cd163*^*-/-*^ mice and explaining only a trending increase in BALF oxidative stress following CFH administration (51). Other studies have highlighted the role of CFH in upregulating pro-inflammatory pathways through TLR4 as well as CFH metabolism and CD163 internalization in upregulating antioxidant and anti-inflammatory pathways, through heme oxygenase 1 (HO-1) and IL-10, respectively (22, 23, 45). However, we did not find that CFH treatment altered *Tlr4* expression in whole lung homogenates, and it equally increased *Hmox1* and *Il10* expression for both WT and *Cd163*^*-/-*^ groups. This could indicate that other cell types in the lung upregulate *Hmox1, Tlr4*, and *Il-10* in response to O_3_, especially as murine *Cd163* is exclusively expressed by tissue resident macrophages in the lung (50). Additionally, these findings could indicate upregulation of compensatory CFH uptake pathways when CD163 is deficient. Cell free heme complexed to hemopexin can bind to CD91 to be internalized (52) and there is evidence for CD163-indpendent engulfment of CFH by alveolar macrophages (51). Future analysis of transcriptional changes targeting resident macrophage populations would prove insightful for understanding effects of CD163-mediated clearance on downstream regulation of antioxidant and inflammatory pathways.

We acknowledge the following limitations in the present study. First, only female mice were used in the experiments reported within this study, which does not allow for examination of sex as a biological variable. Female mice were selected as we have previously reported a sex-dependent effect of O_3_ inhalation which causes increased pulmonary inflammation in female mice (32). Moreover, our initial investigations did not reveal an interaction between sex and *Cd163* expression (data not shown). We also did not observe differences in our human data when stratified by sex. However, we may have not been powered to detect differences with respect to sex, and future studies will focus on powering for sex differences, including effects of luteal and follicular phases, which have effects on circulating iron biomarkers (55). Second, although this study presents evidence for CD163-mediated CFH clearance in the pulmonary response to O_3_, future studies are necessary to establish the source of O_3_-driven CFH accumulation. During O_3_ exposure, increased pulmonary permeability could allow erythrocytes to cross the endothelium into the airspace, where they can release CFH in response to hemolytic stress (56, 57). However, there is also evidence suggesting that type 2 alveolar cells and club cells can produce Hb (58). Third, we acknowledge that there are differences between murine and human Hb, which impact translation of our findings in mice to human outcomes. Not only are there structural differences between murine and human Hb (59), but human Hp also has a higher affinity for Hb than does murine Hp, and murine CD163 is able to directly bind Hp without it being complexed to Hp (60, 61). As a result of these differences, further assessment of CFH clearance in human models is required to better understand its role in pulmonary responses to O_3_.

## Conclusion

The present study describes the protective effects of CD163 following acute O_3_ inhalation, including evidence that supports the novel role of CD163-mediated clearance of CFH in reducing O_3_-induced pulmonary inflammation. This SR is one of many that remains understudied in the context of acute exposure to ambient air pollutants. Moreover, further research is necessary to define other pathways involved in scavenging CFH and hemoglobin derivatives. Cumulatively, these findings indicate that CFH clearance through the CD163 SR may hold therapeutic potential in mitigating O_3_-induced lung injury and inflammation.

## Author contributions

SJC - Conceived and designed research, performed experiments, analyzed data, interpreted results of experiments, prepared figures, drafted manuscript, edited and revised manuscript, approved final version of manuscript.

BS - Conceived and designed research, performed experiments, analyzed data, interpreted results of experiments, prepared figures, edited and revised manuscript, approved final version of manuscript.

ES - performed experiments, analyzed data, edited and revised manuscript, approved final version of manuscript.

KDR - Conceived and designed research, performed experiments, analyzed data, interpreted results of experiments, edited and revised manuscript, approved final version of manuscript.

GMH - performed experiments, edited and revised manuscript, approved final version of manuscript.

AV - performed experiments, analyzed data, interpreted results of experiments, edited and revised manuscript, approved final version of manuscript.

AB - performed experiments, analyzed data, interpreted results of experiments, edited and revised manuscript, approved final version of manuscript.

CR- performed experiments, analyzed data, interpreted results of experiments, edited and revised manuscript, approved final version of manuscript.

TJM - Conceived and designed research, interpreted results of experiments, edited and revised manuscript, approved final version of manuscript.

HZ - performed experiments, analyzed data, interpreted results of experiments, edited and revised manuscript, approved final version of manuscript.

VVK - performed experiments, analyzed data, interpreted results of experiments, edited and revised manuscript, approved final version of manuscript.

MV - performed experiments, analyzed data, interpreted results of experiments, edited and revised manuscript, approved final version of manuscript.

SH – Conceived and designed research, performed experiments, analyzed data, interpreted results of experiments, edited and revised manuscript, approved final version of manuscript.

RMT - Conceived and designed research, interpreted results of experiments, edited and revised manuscript, approved final version of manuscript.

KMG – Conceived and designed research, interpreted results of experiments, edited and revised manuscript, approved final version of manuscript.

