## Supplementary material for "CD163 protects against pulmonary injury and inflammation induced by acute O_3_ exposure": Supplemetal Materials

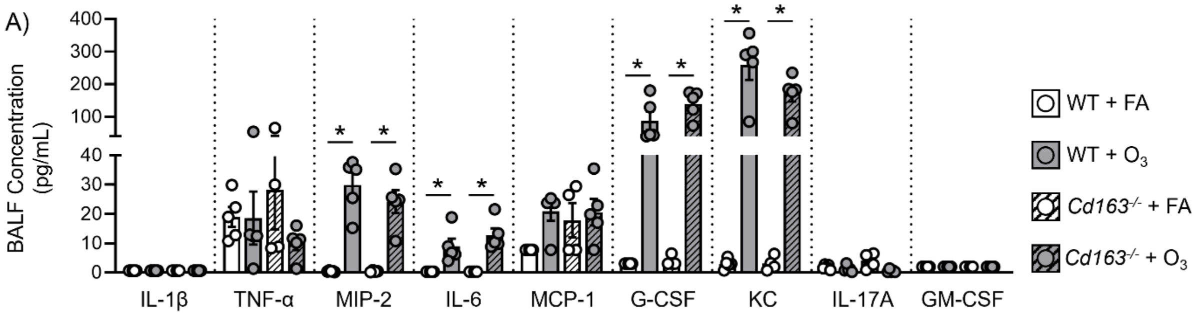


***Supplemental Figure 1. Concentrations of BALF cytokines and chemokines 12 hours after $O_3$ exposure.*** *Female, C57BL/6N wildtype (WT) and $Cd163^{-/-}$ mice, aged 8 – 12 weeks old were exposed to either filtered air (FA) or 1 ppm ozone ($O_3$) for 3 hours. Mice were euthanized 12 and after exposure to collect lung bronchoalveolar lavage fluid (BALF). Multiplex ELISA was used to quantify concentrations of BALF cytokines and chemokines: interleukin-1β (IL-1β); tumor necrosis factor α (TNF-α); macrophage inflammatory protein-2 (MIP-2); interleukin-6 (IL-6); monocyte chemoattractant protein 1 (MCP1); granulocyte colony-stimulating factor (G-CSF); keratinocyte chemoattractant (KC); interleukin-17A (IL-17A), and granulocyte-macrophage colony-stimulating factor (GM-CSF). Mann-Whitney U Test with Holm's adjustment for multiple comparison; n = 4 – 6 /group; *p<0.05*

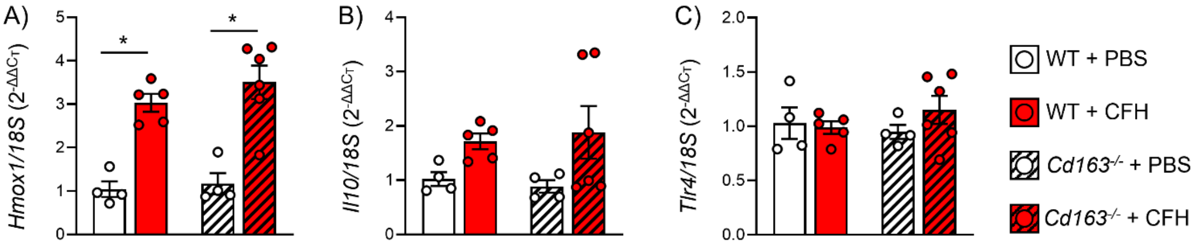


***Supplemental Figure 2. Airspace CFH increases lung tissue Hmox1 and Il10 expression regardless of Cd163 expression.*** *Female, C57BL/6N wildtype (WT) and $Cd163^{-/-}$ mice, aged 8 – 12 weeks old were dosed with 100 µL of either sterile PBS or 1 mg/mL CFH oropharyngeally (o.p.). Mice were euthanized 6 hours to collect lung tissue and bronchoalveolar lavage (BAL) for gene expression, protein analysis, and cell differentials. Lung tissue expression of A) heme oxygenase 1 (Hmox1); B) interleukin 10 (Il10); and C) toll like receptor 4 (Tlr4) were measured using rtPCR. Mann-Whitney U Test with Holm's adjustment for multiple comparison; n = 4 – 6 /group; *p<0.05*
